# Robo1 loss has pleiotropic effects on postnatal development and survival

**DOI:** 10.1101/2023.08.01.551555

**Authors:** Nicole A. Kirk, Ye Eun Hwang, Seul Gi Lee, Kee-Beom Kim, Kwon-Sik Park

**Affiliations:** Department of Microbiology, Immunology, and Cancer Biology, University of Virginia School of Medicine, Charlottesville, VA, USA; Department of Life Sciences, Chung-Ang University, Seoul, KOREA; Department of Life Sciences, Kyungpook National University, Daegu, KOREA

**Keywords:** Robo1, Growth hormone (GH), Insulin-like growth factor 1 (IGF-1), pituitary gland

## Abstract

Roundabout guidance receptor 1 (Robo1) is known to function during midbrain development in mice, but its postnatal role remains poorly defined in part due to the perinatal lethality of the mice lacking the gene. Here, we describe the postnatal phenotypes of *Robo1*^-/-^ mice in the B6/129s genetic background. *Robo1*^*-/-*^ mice had both a slower growth rate and shorter lifespan compared to *Robo1*^+/+^ littermates. Skin histological analysis revealed that *Robo1*^-/-^ mice displayed increased wrinkles, enlarged sebaceous glands, and a reduced subcutaneous fat layer. *Robo1*^*-/-*^ mice displayed accelerated hair graying in line with an observed overall decrease in melanin production. Although both growth hormone (GH) and insulin-like growth hormone 1 (IGF-1) were suppressed in *Robo1*-deficient mice at 3 weeks of age, they were expressed at similar levels to *Robo1* intact mice at 7 weeks of age. These findings suggest that *Robo1*-mediated hormone regulation is required for normal growth during the early stages of puberty and may provide a crucial timepoint at which to examine the molecular mechanisms of pituitary disorders.

## Introduction

Growth and aging are universal processes marked by the progressive accumulation of physical, molecular, cognitive, and physiological changes with time. The pituitary gland produces and secretes vital hormones responsible for proper growth, and almost all of these hormones are altered with aging (Veldhuis, 2013). The embryonic development of the pituitary gland involves a complex and highly regulated network of signaling molecules. The somatotropic axis, including growth hormone (GH) and insulin-like growth factor (IGF-1), is a well-established regulator of postnatal growth in mammals (Aguiar-Oliveira et al., 2019). GH is secreted from the pituitary gland and stimulates the liver to synthesize IGF-1 (Schneider et al., 2003). Their levels peak during puberty to enhance skeletal growth and nutrient acquisition (Poudel et al., 2020). Thus, a growth hormone deficiency results in severe developmental defects caused by either hormone related genetic mutations or defects in the structural developmental of the brain, especially the pituitary gland.

Roundabout guidance receptor 1 (ROBO1) is a member of the immunoglobulin gene superfamily and encodes a transmembrane receptor that plays an important role in neurodevelopment. Specifically, ROBO1 functions to drive axon branching, guide axon pathfinding, and control neuronal migration (Fujiwara et al., 2006; Kidd et al., 1998). Thus, *ROBO1* mutations results in aberrant axonal guidance during brain development including the extension of the growth cone past the midline (Seeger et al., 1993; Andrews et al., 2006; López-Bendito et al., 2007). Because homozygous deletion of *Robo1* is perinatally lethal (Xian et al., 2004a; Andrews et al., 2006), our current understanding of the role of ROBO1 is limited to embryonic development or in the context of a heterozygous knockout (*Robo1*^*+/-*^). It has recently been shown that *ROBO1* mutations are linked to neural diseases including pituitary stalk interruption syndrome (PSIS), combined pituitary hormone deficiency (CPHD), myelodysplastic syndrome, and dyslexia (Bashamboo et al., 2017; Hannula-Jouppi et al., 2005; Xu et al., 2015).

*Robo1* is lost or mutated in various types of cancer, including a subset of small-cell lung cancer (SCLC) (Dallol, 2002; George, 2015). To build a mouse model to study the role of Robo1 loss on SCLC development *in vivo*, we crossed mice with heterozygous *Robo1* null allele (*Robo1*^*+/-*^) with compound transgenic mice with floxed alleles of *Rb, p53*, and *Rbl2* (*Rb*^*lox/lox*^; *p53*^*lox/lox*^; *Rbl2*^*lox/lox*^), which is a mouse model of human SCLC. During the genetic crosses, however, we observed that the mice with *Robo1*^*-/-*^ genotype were born alive and lived well beyond 2 months after birth (Figure 1). This was unexpected since *Robo1* knockout mice are known to die perinatally. The *Robo1* knockout mice displayed several notable defects that were not previously reported, including growth defects and signs of premature aging. This report documents these phenotypes and suggests that *Robo1* is a pleiotropic factor and its reduced function during development has significant implications in postnatal growth and homeostasis.

**Figure 1.**
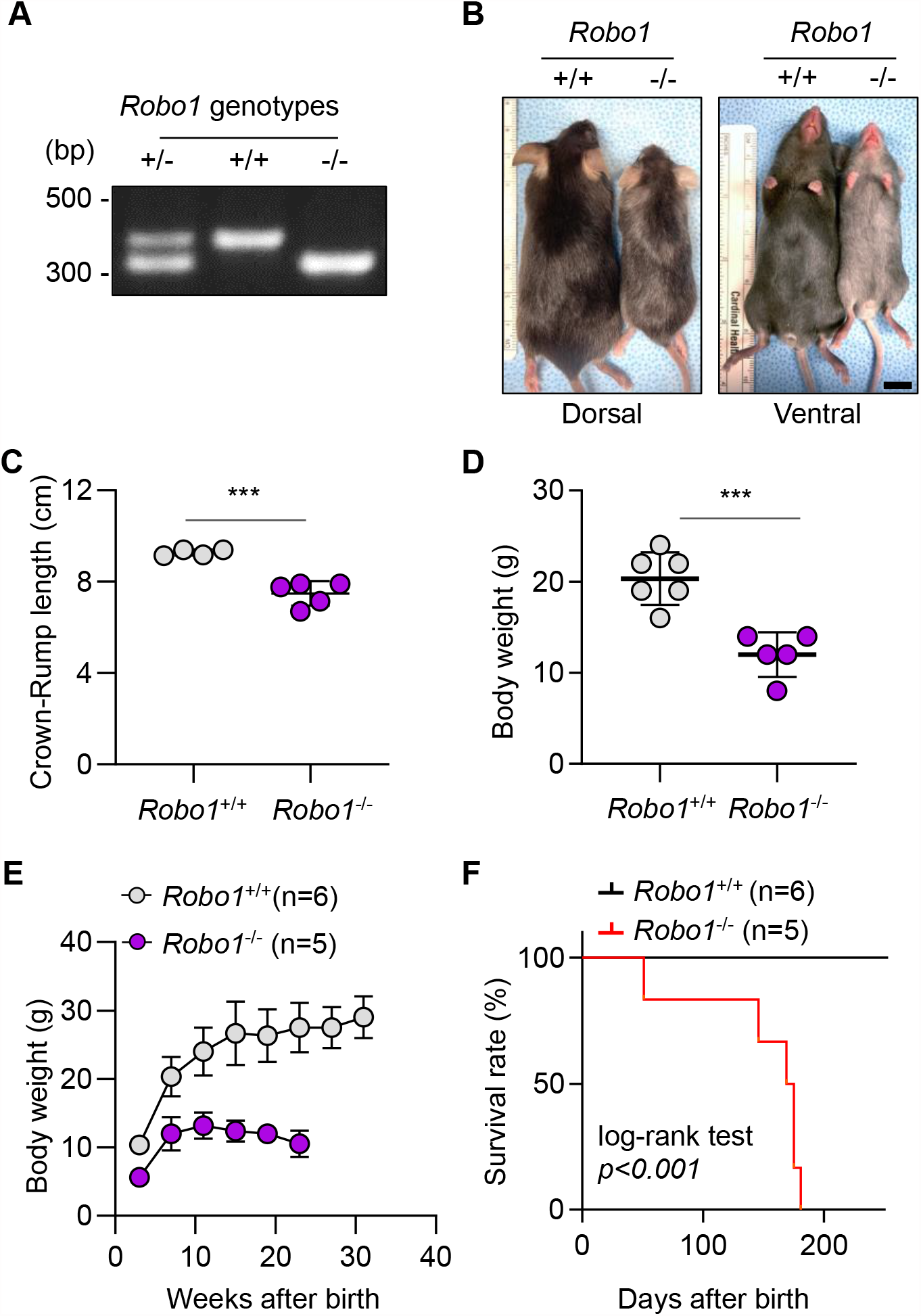
Growth defects and shortened life span in *Robo1* deficient mice. (A) Genotyping results of *Robo1*^+/-^, *Robo1*^+/+^, and *Robo1*^-/-^ mice. (B) Representative images of the dorsal and ventral sides of 7-week-old *Robo1*^+/+^ and *Robo*1^-/-^ mice. Scale bar: 1 cm (C) Crown-rump length and (D) body weight of 7-week-old *Robo1*^+/+^ and *Robo1*^-/-^ mice. (E) Body weight monitored at 4 week intervals starting in 3-week old mice. (F) Kaplan-Meir survival curves of *Robo1*^+/+^ (n=6) and *Robo1*^-/-^ mice (n=5). Log-rank (Mantel-Cox) test was used to determine the significance between *Robo1*^+/+^ and *Robo1*^-/-^ mice. \*\*\**p* < 0.001. Data are presented as the mean ± standard deviation (S.D.). Statical tests were performed using an unpaired t-test.

## Materials and Methods

### Generation of *Robo1*^*-/-*^ mice, genotyping, and monitoring

Mice with heterozygous mutation in *Robo1* (*Robo1*^+/-^) were previously described (Long, 2004). The *Robo1*^+/-^ mice in a pure C56BL/6 (B6) background was kindly provided by the laboratory of Dr. Marc Tassier-Lavigne. *Rb*^*lox/lox*^*;p53*^*lox/lox*^; *Rbl2*^*lox/lox*^ (*RPR2* mice) have been previously described and maintained in a mixed genetic background (B6/1S9S) (Schaffer, 2010). Mice were maintained according to guidelines from the National Institutes of Health. Animal procedures were approved by the Institutional Animal Care and Use Committee at the University of Virginia. We performed genetic crosses between *Robo1*^-/-^ mice and *RPR2* mice and collected tails before weaning of mouse pups. Tails were digested in lysis buffer containing proteinase K (Thermo Fisher Scientific, BP1700-100) and DNAs were purified using potassium acetate and subsequently 50% ethanol. Then, DNAs were resuspended in water and were added to polymerase chain reaction (PCR) mix. Sequences of the primers for *Robo1* genotyping were previously described (Long, 2004), including a forward primer common to both wild-type and mutant alleles 5′-TGGCACGAAGGTATATGTGC-3′, a wild-type allele-specific reverse primer 5′-GAAGGACTGGTGGTTTTGAG-3′, and a mutant allele-specific reverse primer 5′-CCTCCGCAAACTCCTATTTC-3′. The PCR conditions were as follows: 95° for 5 min; 30 cycles of 95° for 20 sec, 58° for 45 sec, and 72° for 45 sec; finally, 72° for 10 min. PCR products were separated in 3% agarose gel and target PCR products were identified under UV light. To measure body length, we measured from the top of the head (crown) to the bottom of the buttock (rump). Mouse growth curves and Kaplan-Mayer survival plots display changes in animal weight at 4-week intervals starting at week 3.

### Histology analysis

Mice were euthanized using an overdose of anesthetic agent as approved by the animal protocol. Various organs and tissues were collected, including the lung, brain, skin and bones, fixed in 4% paraformaldehyde (PFA)/phosphate-buffered saline (PBS) overnight before being processed for paraffin embedding. Five-micron (5 μm) sections of paraffin blocks were stained with hematoxylin and eosin (H&E). To examine melanin deposit in the skin, sections were stained using a Fontana-Masson staining kit (Milipore, HT2000), which is widely used for the histological visualization of melanin in paraffin or frozen sections. Light microscopy images of skin sections stained with H&E or Fontana-Masson were obtained using Nikon Eclipse Ni-U microscope (Nikon). Image analysis and automated quantification of target parameters were performed using NIS-Elements Basic Research software (Nikon).

### Assays for growth hormone (GH) and insulin-like growth factor 1 (IGF-1) in plasma

Littermates of mice were euthanized at 3 and 7 weeks of age using a sub-lethal dose of avertin injection and blood was collected in K2-EDTA tubes (BD Bioscience) using the retro-orbital sinus blood collection method essentially as previously described (Parasuraman, 2010). Plasma was separated using 4° tabletop centrifuge at 1,500g for 15 minutes. The plasma samples were then used for enzyme linked immunosorbent assay (ELISA) to quantify levels of growth hormone (GH) and insulin-like growth factor 1 (IGF-1) using commercial ELISA kits (AFG Scientific, EK730389 and Proteintech, KE10032, respectively) according to the manufacturers’ instructions. Plasma was diluted in sample diluent (1% bovine serum albumin, 0.05% Tween-20 in PBS) at 1:5 for GH ELISA and 1:300 for IGF-1 ELISA and stored at -20° until the assay was performed. Processed samples in the ELISA were read at 450 nm/650 nm using spectrophotometer (BioTek).

### Statistical analysis

Statistical analysis and drawing graphs was performed with GraphPad Prism 8.0 (GraphPad Software). Kaplan-Meier curves were used for plotting mice survival and log-rank test was used to determine significance with *p*<0.05 considered to be significant. One-way Analysis of Variance (ANOVA) was used for comparisons of more than two groups. One-way ANOVA F values were displayed in each figure legend as F (D_Fn_, D_fd_) (D_Fn_ as the df nominator and D_fd_ as the df denominator). Tukey’s multiple comparison test was applied to all ANOVA analyses post-hoc test. Unpaired Student t-tests were used for the comparison of two groups. Results are presented as the mean ± standard deviation (S.D.) and statistical significance was set as *p* <0.05 (\**p*<0.05, \*\**p*<0.01, and \*\*\**p*<0.001).

## Results

### *Robo1*-deficient mice have growth defects and a shortened life span

We examined the genotypes of newborns from the cross of *Robo1*^*+/-*^ male and females using genotyping PCR and confirmed that each litter generally has 1:2:1 ratio of *Robo1* wild type, heterozygous, or homozygous mutant genotypes (Figure 1A and not shown). This was unexpected given the previous reports showing that complete deletion of *Robo1* was perinatally lethal in B6 mice (Xian et al., 2004a; Andrews et al., 2006). However, we noted visible physical differences including distinct growth alterations in body size and weight between the *Robo1*^+/+^ and *Robo1*^-/-^ mice (Figure 1B, C). At seven weeks of age, body length (the crown rump length) and weight of *Robo1*^-/-^ mice was significantly smaller compared to *Robo1*^*+/+*^ mice (Figure 1C, D). To determine whether the growth was simply delayed in *Robo1*-deficient mice, we monitored several cohorts of mice over a period of time with 4-week intervals starting when they were 3-week old. The period of rapid growth in both *Robo1*-intact and deficient mice is typically observed between weeks 3 through 7, the established timeframe of mouse puberty. At 3 weeks of age, both the weight and overall growth rate of *Robo1*^-/-^ was significantly lower than *Robo1*^+/+^ mice and the degree of separation between the weight of *Robo1*^*-/-*^ and *Robo1*^*+/+*^ mice increased over time (Figure 1E). Although *Robo1*^-/-^ survived through the period of growth in normal mice, Kaplan-Meyer survival curve showed that the median survival age of *Robo1*^-/-^ mice was 172 days, significantly shorter than that of *Robo1*^+/+^ mice (Figure 1F). *Robo1*^-/-^ mice gradually became less active and their movement slowed as the mice reached 2 months of age whereas *Robo1*^*+/+*^ littermates become more active and faster as expected during this time period (not shown). These phenotypes of *Robo1* were observed in both male and female mice (Figure 2A). Together, these findings suggest that depletion of *Robo1* results in growth defects and a shortened life span in mice.

**Figure 2.**
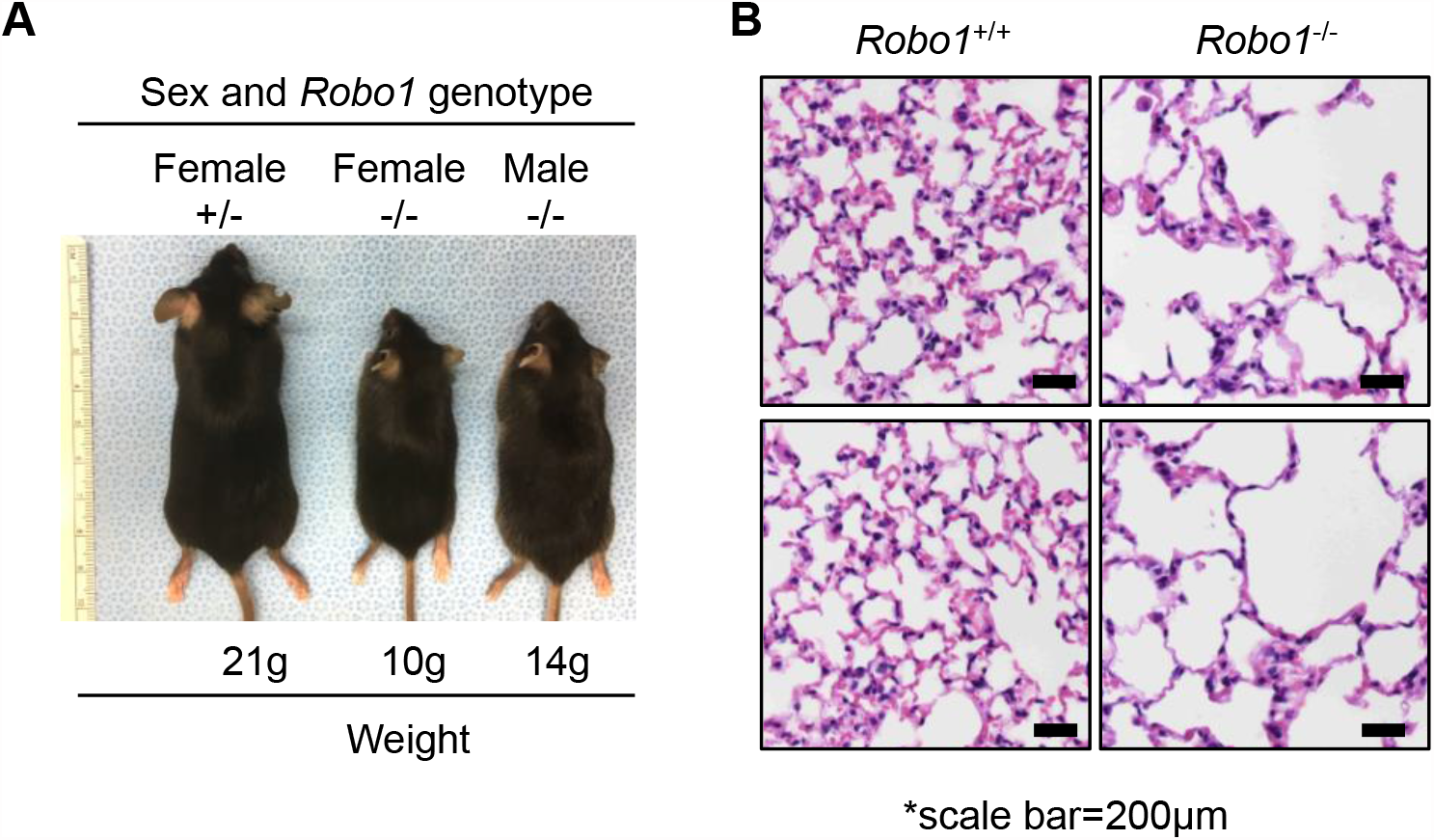
Impact of Robo1 deficiency on both male and female mice and the lung alveoli. (A)The growth defects in *Robo1*^-/-^ mice affect both female and male mice. (B) Representative images of H&E stained sections of peripheral lungs. Alveolar spaces are enlarged in *Robo1*^-/-^ deficient lungs relative to control littermates.

### *Abnormalities in skin of Robo1*-deficient mice

The abnormal body size and weight were not the only atypical phenotypes observed in *Robo1* deficient mice. We observed enlarged alveolar spaces in the *Robo1* deficient lungs relative to control littermates (Figure 2B), which was previously reported (Branchfield, 2016). Other remarkable phenotypes include the presence of gray skin hairs at a very early age (Figures 1B and 3A). The hair graying was more pronounced on the ventral skin rather than on the dorsal skin (Figures 3A). To determine an origin of hair graying at the tissue level, we examined melanin using Fontana-Masson staining (Sheehan and Hrapchak, 1973). The staining confirmed less melanin deposits in the hairs in *Robo1*^-/-^ mice relative to *Robo1*^*+/+*^ littermate controls (Figure 3A). Although we did not detect melanin staining in the epidermal layer of the skin in neither group of mice, the melanin stain pattern suggests that hair greying was the result of reduced melanin. We also noted aberrant histology of the skin layer in *Robo1*^-/-^ mice. H&E staining on ventral skin tissues in both mice showed increased wrinkles and reduced subcutaneous fat layers in *Robo1*^-/-^ mice relative to *Robo1*^*+/+*^ littermate controls (Figure 3B, C). Additionally, the sebaceous glands in in hair follicles were significantly enlarged in in *Robo1*^-/-^ mice relative to *Robo1*^*+/+*^ controls. (Figure 3B). Taken together, these results suggest that the loss of *Robo1* results in alterations in the skin that are apparent through histological examination.

**Figure 3.**
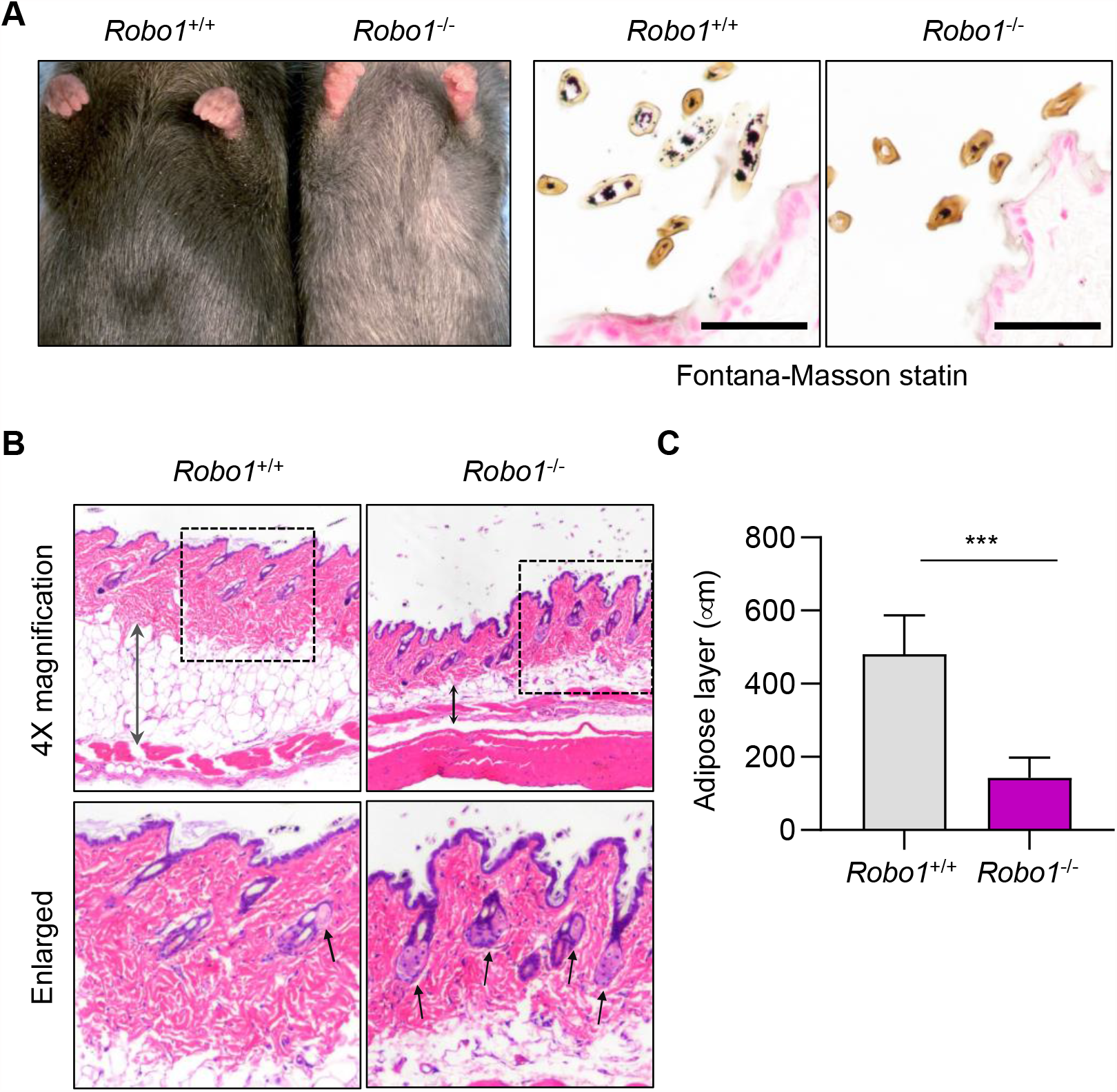
Abnormalities in skin of Robo1-deficient mice. (A) Representative picture of ventral skins. The hairs in *Robo1*-deficient mice grayer more than those in control litter mates. (B) H&E-stained skin section from *Robo1*^+/+^ and *Robo1*^-/-^ mice. Black arrows indicate sebaceous glands, and the double-headed black arrows indicate thickness of the adipose layers. (C) Plot of the adipose layer thickness in *Robo1*^+/+^ and *Robo1*^-/-^ mice. Results are average of 3 lines per section, and 5 sections per mouse (n=3 mice per group). (Data are presented as the mean ± S.D.. Statical tests were performed using unpaired t-test.

### Reduced levels of growth hormone (GH) and insulin-like growth factor 1 (IGF-1) in the plasma of *Robo1*-deficient mice

An axis of growth related hormones, such as GH and IGF-1, has been established as a key regulator of development and mammalian growth (Dodds and Noble, 1936). The pituitary gland is a major endocrine gland that regulates homeostasis through secretion of various regulatory hormones, including GH. Since the average size of the pituitary gland of *Robo1* mutant patients was small (Bar et al., 2015; Bashamboo et al., 2017; Liu and Chen, 2020, we postulated that the short stature in *Robo1*^-/-^ mice is linked to possible defects in growth hormone release by the pituitary gland. To demonstrate the difference of GH level between *Robo1*^+/+^ and Robo1^-/-^ mice in case of 3-weeks and 7-weeks old, we measured GH concentration in mice plasma using ELISA. At 3-weeks old, the GH levels were significantly reduced in *Robo1*^-/-^ mice compared to *Robo1*^*+/+*^ mice while there was no difference once they reached 7-weeks of age (Figure 4B). GH impacts growth and aging directly after secretion by the pituitary gland and also indirectly by stimulating the synthesis and release of IGF-1 into the bloodstream by the liver (Carter et al., 2002). Similar to the pattern observed in GH levels (Figure 4B), the 3-week old *Robo1*^*-/-*^ mice had reduced IGF-1 levels compared to their *Robo1*^*+/+*^ counterparts but this difference was lost once the mice reached 7-weeks (Figure 4B). Collectively, these findings suggest that the decrease in the size of the pituitary gland results in suppression of the levels of GH and IGF1 in *Robo1*^-/-^ mouse plasma (Figure 4C).

**Figure 4.**
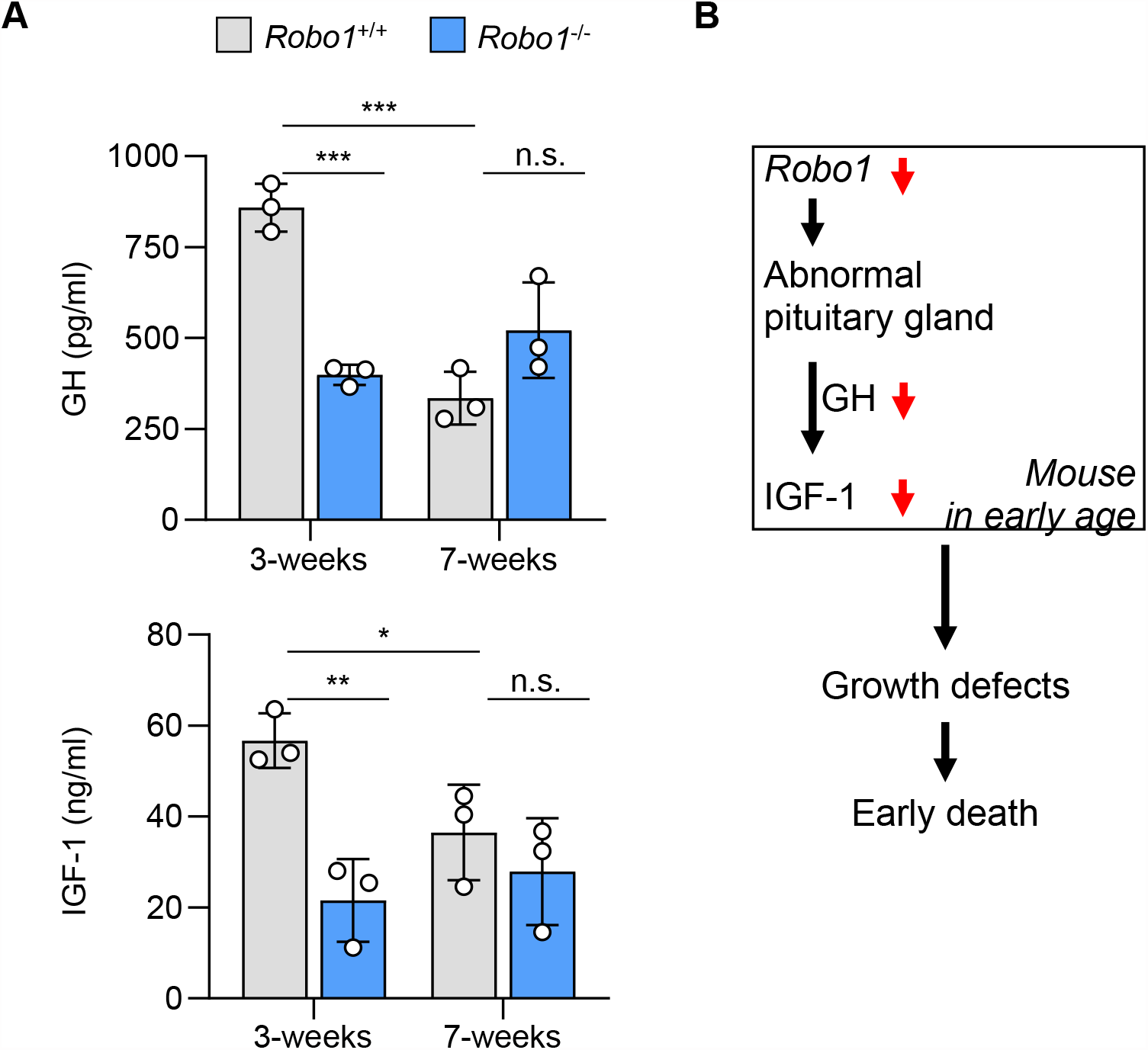
Reduced levels of GH and IGF-1 in the plasma of *Robo1*-deficient mice. (A) Top, GH concentration in *Robo1*^+/+^ and *Robo1*^-/-^ plasma. One-way ANOVA, F_3,8_=23.50. Bottom, IGF-1 concentration in *Robo1*^+/+^ and *Robo1*^-/-^ plasma. One-way ANOVA, F_3,8_=7.63. (B) Model for the role of the Robo1-mediated GH/IGF1 axis on regulating mouse pubertal growth. Data is presented as the mean ± S.D. from three independent experiments. \**p* < 0.05, \*\**p* < 0.01, \*\*\**p* < 0.001, n.s. not significant.

## Discussion

The role of *Robo1* in postnatal development remained largely unknown due to the perinatal lethality of complete *Robo1* loss in mice. In this study, we were able to generate homozygous *Robo1* knockout mice that lived longer than 3 moths of age. It is possible that the mixed genetic background (B6/129S) of these mice contributed to the extended survival of these mice through compensation for *Robo1* loss, which does not occur in mice with a pure B6 background (Xian, 2004b). This new model of *Robo1* deficiency revealed various phenotypes. Most notably, *Robo1* deficient mice were born with significant growth defects and died prematurely around the reproductive age. Accompanying phenotypes include gradual loss of movement, enlarged alveolar spaces in the lung, and abnormality in the skin featuring increased wrinkles, hair graying, and reduced subcutaneous fat. In line with the growth defects, plasma levels of GH and IGF-1 in the *Robo1* deficient mice at the time of weaning were significantly lower than *Robo1* wild type littermates. These findings suggest that the germline loss of *Robo1* has pleiotropic effects on postnatal development ranging from growth to life expectancy.

The underlying mechanisms of the pleiotropic effects are likely diverse. Our results imply that the reduced levels of systemic GH and IGF-1 underlie the growth defects observed in the *Robo1* deficient mice as the GH/IGF-1 axis regulates postnatal growth in mammals, especially during puberty (Lupu et al., 2001; MacKelvie et al., 2002; Stratikopoulos et al., 2008). Notably, the concurrent decreases in GH and IGF1 that were present at the time of weaning 3 weeks after birth but were absent at 7-weeks of age corresponding to the late part of the pubertal growth (Dutta and Sengupta, 2016), suggest that the activity of these hormones before the onset of puberty is crucial. It has been reported that GH and IGF-1 affect not only pubertal growth, but also skin organization (Kanaka-Gantenbein et al., 2016). Excessive GH levels can lead to acromegaly, neurofibromatosis-1 (NF-1) and carney complex (CNC), all of which are characterized by increased melanin and skin pigmentation (Corcuff et al., 1997; Landau and Krafchik, 1999; Pack et al., 2000). Conversely, a growth hormone deficiency (GHD) is linked with thin and dry skin, increased wrinkles, and irregular sweating (Conte et al., 2000). Taken together, the similarities between the phenotypes of *Robo1* deficient mice and the symptoms of human growth hormone-related disorders further suggest that the growth defects and skin abnormalities in *Robo1* deficient mice are attributed to the hyposecretion of GH and IGF-1.

The phenotypes of *Robo1* deficient mice may be clinically relevant. *Robo1* mutations are associated with pituitary stalk interruption syndrome (PSIS) and combined pituitary hormone deficiency (CPHD) in humans (Bar et al., 2015; Bashamboo et al., 2017; Liu and Chen, 2020). These congenital disorders manifest as growth hormone deficiencies, lower body weights, and delayed growths in height, which is caused by pituitary dwarfism and the resulting deficiency in pituitary hormone release. While the underlying molecular mechanisms of these disorders remain unknown, it has been suggested that mutations in *POU1F1, PROP1*, and *PTCH1* contribute to PSIS (Brauner et al., 2020; Diwaker et al., 2022). For the first time, our study provides functional evidence for the role of *Robo1* deficiency as a cause for human disorders and presents a promising potential animal model.

In conclusion, we identified a novel role for *Robo1* in somatotrophic growth. The ablation of *Robo1* led to a malformation of the pituitary structure during early puberty and defects in secretions of pituitary related hormones which resulted in a lower growth rate aberrant skin histology. The study provides novel insights into hormone regulation and further demonstrates the potential for *Robo1*^*-/-*^ mice to be utilized in studying the molecular mechanisms underlying PSIS.

